# pyCapsid: Identifying dominant dynamics and quasi-rigid mechanical units in protein shells

**DOI:** 10.1101/2023.02.27.529640

**Authors:** Colin Brown, Anuradha Agarwal, Antoni Luque

## Abstract

**Summary:** pyCapsid is a Python package developed to facilitate the characterization of the dynamics and mechanical units of protein shells and other protein complexes. The package was developed in response to the rapid increase of high-resolution structures, particularly capsids of viruses, requiring multiscale biophysical analyses. Given a protein shell, pyCapsid generates the collective vibrations of its amino-acid residues, identifies quasi-rigid mechanical regions, and maps the results back to the input proteins for interpretation. pyCapsid summarizes the main results in a report that includes publication-quality figures.

**Availability and Implementation:** pyCapsid’s source code is available under MIT License on GitHub (https://github.com/luquelab/pycapsid). It is compatible with Python 3.8-3.10 and has been deployed in two leading Python package-management systems, PIP (https://pypi.org/project/pyCapsid/) and Conda (https://anaconda.org/luque_lab/pycapsid). Installation instructions and tutorials are available in the GitHub Page-style online documentation (https://luquelab.github.io/pyCapsid). Additionally, a cloud-based implementation of pyCapsid is available as a Google Colab notebook (https://colab.research.google.com/github/luquelab/pyCapsid/blob/main/notebooks/pyCapsid_colab_notebook.ipynb). pyCapsid Colab does not require installation and generates the same report and outputs as the installable version. Users can post issues regarding pyCapsid in the GitHub repository (https://github.com/luquelab/pyCapsid/issues).

## 1 Introduction

Viruses protect their infective genomes in protein shells called capsids (Twarock and Luque 2019). The number of capsid structures solved at high-resolution has increased exponentially in the last two decades, partly thanks to cryo-electron microscopy advances (Callaway 2020; Johnson and Olson 2021; Montiel-Garcia et al. 2021). These three-dimensional reconstructions combined with computational algorithms and complementary experimental techniques are leading to a mechanistic characterization of the assembly, dynamics, and stability of viral capsids, opening the doors to new antiviral strategies (Johnson et al. 2021; Twarock and Stockley 2019; Organtini et al. 2017; Kizziah, Rodenburg, and Dokland 2020; Yeager et al. 1990; Qazi et al. 2018; Li et al. 2008; Mata et al. 2020; de Pablo and San Martín 2022; X. Zhang et al. 2013; Bayfield, Steven, and Antson 2020; Montiel-Garcia et al. 2021; Podgorski et al. 2020; Hua et al. 2017; Mohajerani et al. 2022; Wilson and Roof 2021; Bruinsma, Wuite, and Roos 2021; Luque and Reguera 2013; Lee et al. 2022; Grime et al. 2016; Plavec et al. 2021). Among the computational methods, molecular dynamics algorithms have improved dramatically in the last decades and can infer the dynamics of large protein complexes. However, they resolve relatively short timescales (⪝1 μs) and require specialized computational resources (Jana and May 2021; Hadden et al. 2018; Perilla and Schulten 2017; Bryer et al. 2022). The fact that capsids are assembled from 60 to more than 60,000 proteins further limits the application of molecular dynamics (Twarock and Luque 2019; Luque et al. 2020; Berg and Roux 2021). Alternatively, the combination of normal mode analysis (NMA), molecular coarse-graining, and elastic network models (ENM) offers a more scalable solution (Bahar et al. 2010; Romo and Grossfield 2011). This approach has successfully estimated the collective motion of proteins in complexes (Bahar et al. 2010) or identified structural conformational changes in capsids (Tama and Brooks 2005). Nonetheless, no easy-to-use computational packages are currently available to characterize the dynamics and mechanical properties of protein shells. The bioinformatics software presented here, pyCapsid, aims to address this issue.

pyCapsid is inspired by prior publications that applied ENM, NMA, and clustering methods to extract the quasi-rigid regions of protein shells (Ponzoni et al. 2015; Polles et al. 2013). These methods can identify the mechanical units involved in the assembly or disassembly of protein shells (Polles et al. 2013). However, obtaining these results using packages such as PRINQ++ (Polles et al. 2013), SPECTRUS (Ponzoni et al. 2015), NRGTEN (Mailhot and Najmanovich 2021), ClustENMD (Kaynak et al. 2021), WebPSN (Seeber et al. 2015), or the popular ProDy (S. Zhang et al. 2021), is not trivial and limited to relatively small complexes. To address this issue, we introduce pyCapsid as an accessible and efficient Python package that identifies the dominant dynamics and quasi-rigid regions of protein shells. The underlying methods used in pyCapsid are generic and can be applied to other protein complexes, yet, in this first release of pyCapsid, we have focused on the characterization of protein shells, such as viral capsids, cellular protein compartments like encapsulins, and gene-transfer agents (Giessen et al. 2019; Bárdy et al. 2020; Johnson and Olson 2021; Montiel-Garcia et al. 2021).

## 2 Methods and Features

pyCapsid’s Python package is divided into five independent modules (Figure 1): The PDB (protein data bank) module, the CG (coarse-graining) module, the NMA (normal mode analysis) module, the QRC (quasi-rigid clustering) module, and the VIS (visualization) module. The role and technical aspects of each module are briefly described below.

**Figure 1.**
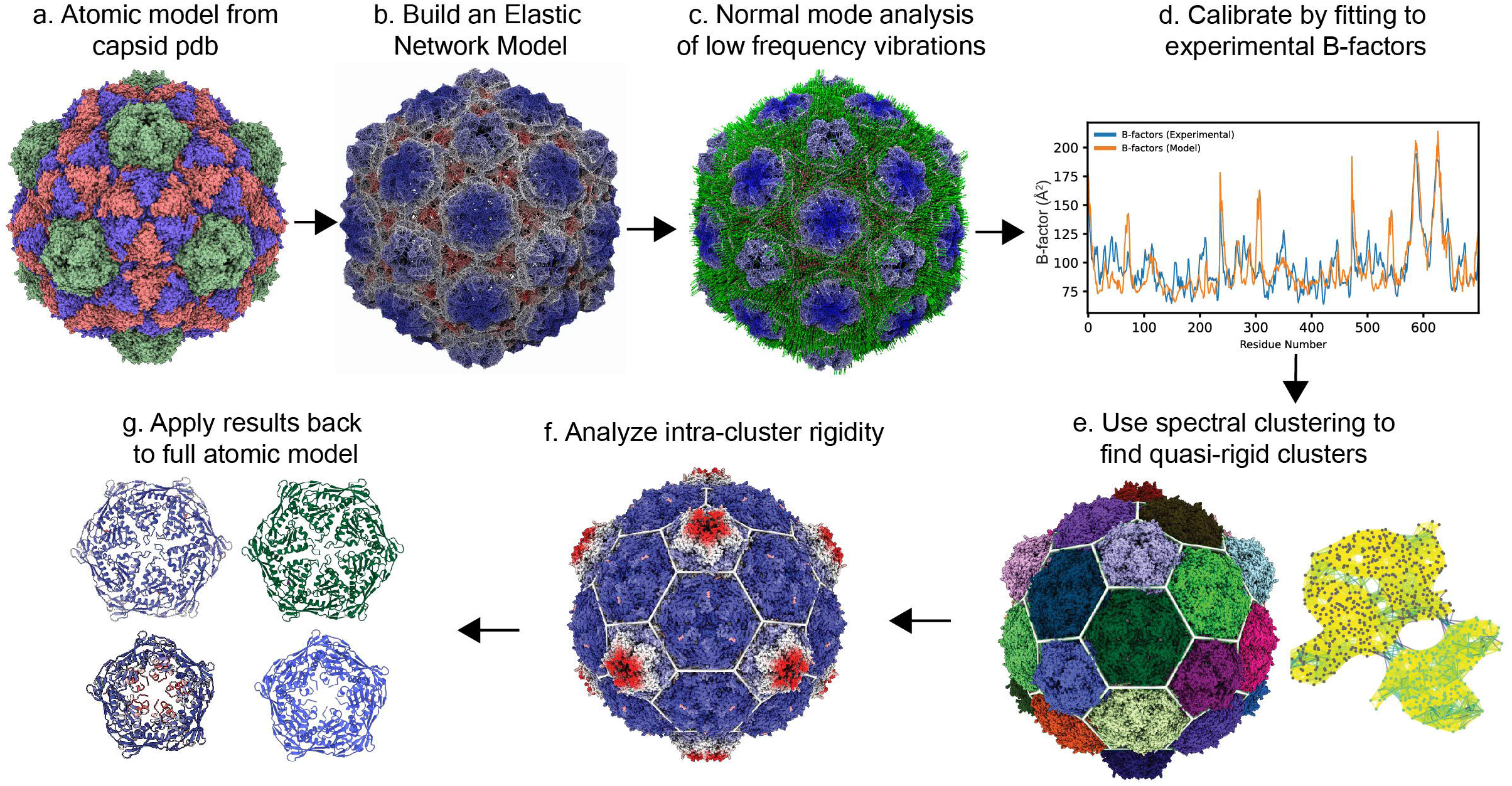
pyCapsid’s key steps. a. The protein shell (atom coordinates, atom types, and B-factors) is inputted using a PDB file (or a PDB ID). b. The elastic network model (ENM) is calibrated (parametrized). c. Normal mode analysis (NMA) determines the distance fluctuations between residues. d. The correlation coefficient of predicted and experimental B-factors is generated for quality control. e. Spectral clustering is applied to obtain the quasi-rigid molecular domains in the capsid. f. The fluctuations between residues within the rigid clusters are calculated. g. The results are mapped back to the capsid protein atomic model for structural interpretation.

### PDB module

This module retrieves and loads structural data from the Protein Data Bank (using the PDB ID) or a local file in PDB or PDBx/mmCIF formats (Westbrook et al. 2022; Berman et al. 2000). The PDB module builds on functions from the Python package Biotite (Kunzmann and Hamacher 2018).

### CG module

This module coarse-grains the proteins at the amino-acid level and establishes an elastic force field between amino acids. It offers four different elastic models: The anisotropic network model (ANM), the Gaussian network model (GNM), the generalized anisotropic or unified elastic network model (G-ANM or U-ENM), which is the default, and the backbone-enhanced elastic network model (bb-ENM). Each amino acid is coarse-grained as a point mass on the alpha-carbon. The links in the network connect amino acids that are within a threshold. The default value is 15Å for ANM and 7.5Å for the other models. These values are based on prior studies of elastic models reproducing empirical molecular thermal fluctuations (B-factors) (Zheng 2008; Micheletti, Carloni, and Maritan 2004; Eyal, Yang, and Bahar 2006; Romo and Grossfield 2011). The small threshold distance leads to a sparse network. This network, combined with the elastic strength values of the elastic model, defines the Hessian matrix. The calculations to build the matrix are accelerated using Numba (Lam, Pitrou, and Seibert 2015).

### NMA module

This module obtains the motions of the macromolecular complex by decomposing the dynamics into independent sinusoidal motions called normal modes (Goldstein, Poole, and Safko 2002). The normal modes and associated frequencies are obtained from the Hessian matrix derived in the CG module. It is well established that only low-frequency modes are relevant to the global dynamics of macromolecules (Bahar et al. 2010). The default number of modes calculated in pyCapsid is 200. This number was selected by comparing the results with simulations using a larger number of modes (as many modes as 1/100 of the number of residues, that is, 1000 modes for a structure containing 100,000 residues). pyCapsid also provides an optional dependency to accelerate the calculations in GPUs using CUDA via solvers in the cupy package (Okuta et al. 2017).

### QRC module

This module estimates the amino acids that tend to fluctuate as a single mechanical unit (quasi-rigid cluster) using the SPECTRUS algorithm (Ponzoni et al. 2015). A cluster contains groups of residues that minimize the distance fluctuations between residues. The clustering methods in pyCapsid are the default discretize method from scikit-learn (Pedregosa et al. 2012), and k-means clustering is offered as an alternative (Yu and Shi 2003). pyCapsid explores a range of clusters from four to a maximum number (n_cluster_max) set by the user. The quality score defined by (Ponzoni et al. 2015) is calculated for each number of clusters, and the set of clusters with the maximum score is selected. If multiple local maxima are observed, each can be selected via the API for further analysis. These alternative maxima can also be obtained by restricting the number of clusters when running pyCapsid.

### VIS module

The results obtained from pyCapsid are stored and organized in data files and figures in the same running folder. pyCapsids’ online tutorial (https://luquelab.github.io/pyCapsid/tutorial/) provides instructions and scripts to visualize the results using the molecular visualization tools NGLview and ChimeraX (Nguyen, Case, and Rose 2018; Pettersen et al. 2021). The results from the cloud-based Colab notebook include a script that can be run directly in Chimera X to generate the 3D visualizations. To obtain high-quality visualizations and animations when running pyCapsid locally, users must have installed ChimeraX (version 1.5 or above) and indicate the path in pyCapsid’s configuration file.

### Execution

Users running the pyCapsid locally can specify the necessary parameters in a configuration file (TOML format), run it, and obtain the results. Users running pyCapsid on the cloud as a Colab notebook can modify parameters as indicated in the notebook’s quick-start guide. Providing the pyCapsid Colab option follows the accessibility trend of cloud-based bioinformatics tools in this environment, like ColabFold (Mirdita et al. 2022) and other tools predicting protein complexes (Bryant et al. 2022). For advanced users, pyCapsid provides an API to access the objects in the five separate modules described above.

## 3 Applications

### Performance and accuracy

pyCapsid’s performance and accuracy were obtained by studying 25 protein shells, which contained from 16,000 to 400,000 amino acid residues, displayed icosahedral symmetry with T-numbers spanning from T=1 to T=16, and had resolutions ranging from 2Å to 5.2 Å. The baseline performance was obtained using an HPC cluster core with Intel Xeon CPU E5-2650 v4 (2.20 GHz) and 128 GB of RAM. The peak memory usage ranged from 800 MB to 90 GB and increased with the number of residues following a power law (exponent = 1.46±0.06 and R^2^ = 0.97). The runtime ranged from 2 minutes to 36 hours and increased with the number of residues following a power law (exponent = 2.20±0.10 and R^2^ = 0.95). The correlation coefficient between the simulated and empirical thermal motions (B-factors) of the amino acids was used as a proxy to evaluate the accuracy of the selected elastic network model (ENM) and generated normal mode analysis (NMA). The correlation coefficients ranged from 0.10 to 0.88 out of 1.00. The distribution of correlation coefficients was consistent with the correlations observed for B-factors predicted using ENM and NMA in smaller protein complexes (Eyal, Yang, and Bahar 2006). The accuracy decreased linearly for structures with lower experimental resolution (slope = –0.20 ± 0.05 1/Å and R^2^ = 0.40), with a regression projecting perfect accuracy for structures with an ideal experimental resolution of 0 Å (intercept = 1.23 ± 0.18). The accuracy was independent of the number of residues (Spearman’s coefficient = –0.09 and p-value = 0.66). pyCapsid’s performance was also assessed by analyzing five small-to-medium capsids in a free cloud-based Colab account and three personal computers with different configurations. The execution times were on the same order of magnitude as in the HPC, with average relative runtime factors ranging from 0.98 (faster) to 2.66 (slower). The main bottleneck was memory, which placed an upper limit to the largest capsid size that could be analyzed. See SI for further details.

### Benchmarking

The five smallest protein shells were used to benchmark the speed and accuracy of pyCapsid with respect to ProDy. Since ProDy does not generate the quasi-rigid domain decomposition, the comparison focused on the modules responsible for loading the PDB and generating the normal modes analysis (NMA). The anisotropic network model (ANM) was available in both ProDy and pyCapsid and yielded the same B-factors. The unified elastic network model (G-ANM or U-ENM), which is available in pyCapsid but not ProDy, improved by five times the average correlation coefficient (from 0.11 for ANM to 0.56 for U-ENM) of the B-factors using the default number of modes. pyCapsid displayed an average speed increase of 3.0±1.5 with respect to ProDy. This increase was independent of capsid size (Spearman’s coefficient = 0.11 and p-value = 0.76). The increase in speed was due to the use of Numba and the invert shift mode in SciPy. This, however, caused a similar increase in memory usage. In any case, ANM or U-ENM did not impact pyCapsid’s speed performance when using the same number of modes. Thus, U-ENM was selected as the default and recommended model in the pyCapsid package. pyCapsid was not benchmarked quantitatively with tools other than ProDy because either even the smallest protein shells exceeded their capacity, or we encountered technical barriers when deploying them locally to analyze such protein shells. Nonetheless, four additional small capsids (PDB IDs 2ms2, 1za7, 1a34, and 3nap) were investigated by pyCapsid recovering the quasi-rigid domains published previously with PISQRD++ (Polles et al. 2013) and SPECTRUS (Ponzoni et al. 2015). See SI for further details.

### Capsid disassembly

An immediate application of pyCapsid is the ability to identify the capsid regions and protein clusters that are most likely to be involved in the initial capsid disassembly pathway, which is crucial for the development of antiviral strategies. A current experimental issue is the fact that capsids are very stable, so generic stressors and denaturants are used to accelerate the disassembly to make studies *in vitro* possible (Zhao et al. 2012; Zhou, Zlotnick, and Jacobson 2022). pyCapsid’s predictions identify molecular targets that will guide more mechanistic and capsid-specific disassembly experiments to help bridge *in vivo* and *in vitro* conditions. The mechanical units predicted by pyCapsid can be tested experimentally, for example, using atomic force microscopy (Martín-González et al. 2023; Ortega-Esteban et al. 2013), mass spectrometry (Uetrecht et al. 2011; Bond et al. 2020), and light scattering (Garmann, Goldfain, and Manoharan 2019; Timmermans et al. 2022).

## 4. Concluding Remarks

pyCapsid can generate the collective motion and extract the quasi-rigid functional regions of protein shells and other protein complexes. The underlying algorithm of pyCapsid generates the dynamical modes faster than established protein dynamics packages and can handle the quasi-rigid domain decomposition of large protein complexes in the range of minutes to over a day, even in regular computers. The computational efficiency of pyCapsid, combined with its accessibility via Python distribution packages, Google Colab, and online tutorials, will benefit researchers in physical virology, structural bioinformatics, and related fields and will facilitate the prediction of disassembly units in protein shells, fostering new antiviral and drug delivery strategies.

## Acknowledgments

The authors thank the insights from the SDSU Biomath Group, in particular, professors Arlette Baljon and Parag Katira. The authors also thank the SDSU Computational Research Science Center for the HPC used to test and benchmark pyCapsid.

